# Application of personalized differential expression analysis in human cancer proteome

**DOI:** 10.1101/2021.07.18.452812

**Authors:** Liu Yachen, Lin Yalan, Wu Yujuan, Zhang Zheyang, Tong Mengsha, Yu Rongshan

## Abstract

Owing to the recent technological advances, liquid chromatography-mass spectrometry (LC-MS)-based quantitative proteomics can measure expression of thousands of proteins from biological specimens. Currently, several studies have used the LC-MS-based proteomics to measure protein expression levels in human cancer. Identifying differentially expressed proteins (DEPs) between tumors and normal controls is a common way to investigate carcinogenesis mechanisms. However, most statistical methods used for DEPs analysis can only identify deregulated proteins at the population-level and ignore the heterogeneous differential expression of proteins in individual patients. Thus, to identify patient-specific molecular defects for personalized medicine, it is necessary to perform personalized differential analysis at the scale of a single sample. To date, there is a scarcity of systematic and easy-to-handle tool that could be used to evaluate the performance of individualized difference expression analysis algorithms in human cancer proteome. Herein, we developed a user-friendly tool kit, IDEP, to enable implementation and evaluation of personalized difference expression analysis algorithms. IDEP evaluates five rank-based tools (RankComp v1/v2, PENDA, Peng and Quantile) through classic computational and functional criteria in lung, gastric and liver cancer proteome. The results show that the within-sample relative expression orderings (REOs) of protein pairs in normal tissues were highly stable, which provided the basis for individual level DEPs analysis. Moreover, these individualized difference analysis tools could reach much higher efficiency in detecting sample-specific deregulated proteins than the group-based methods. Pathway enrichment and survival analysis results were dataset and analysis method dependent. In summary, IDEP has integrated necessary toolkits for individualized identification of DEPs and supported flexible methods evaluation analysis and visualization modules. It could provide a robust and scalable framework to extract personalized deregulation patterns and could also be used for the discovery of prognostic biomarkers for personalized medicine.

## Introduction

Owing to the rapid development of liquid chromatography-mass spectrometry (LC-MS)-based quantitative proteomics, we can now measure thousands of proteins in biological specimens. The Clinical Proteomic Tumour Analysis Consortium (CPTAC) was launched in 2006 to develop LC-MS-based proteomic measurements in human tumor tissues^[1]^. They have applied its standardized workflows in 2011 to three tumor types (colorectal, ovarian, and breast) ^[2–4]^ from The Cancer Genome Atlas (TCGA).Then, the proteogenomics landscape of 13 other kinds of cancer types have been comprehensively characterized. Key bioinformatic analysis and insights from these CPTAC studies included exploring the trans-acting genomic aberrations on protein expression, re-classifying molecular subtypes based on proteomics, and identifying pathways using Phosphoproteomics^[1]^. However, one important issue of published CPTAC studies is that they profiled few normal tissues. The heterogeneity introduced by variation from different test subjects can further complicate proteomics analyses and limit the ability to profile individually altered cancer proteome. Currently, several studies have used the LC-MS-based proteomics to focus on the measurement of proteins in paired tumor and adjacent tissues including hepatocellular carcinoma^[5,6]^, lung adenocarcinoma (LUAD)^[7–9]^ and gastric cancer^[10]^.These studies provide valuable resources that significantly expand the knowledge of human cancer biology and a more comprehensive picture linking cancer ‘‘genotype’’ to ‘‘phenotype’’ through functional proteomics.

Identifying differentially expressed proteins between tumors and normal controls is a common way to investigate carcinogenesis mechanisms. Most of the studies mentioned above identified the cancer-related proteins using T-test, Significance Analysis of Microarrays (SAM)^[11]^ and Wilcoxon signed-rank test, which are designed to detect the population-level differentially expressed genes (DEGs). The aim of population differential analysis is to detect consistently up or down regulated genes, in average, which do not provide precise information at the individual level and could mask the heterogeneity of differential expression in individuals. Taking the heterogeneous nature of disease into account, several studies have reported outlier detection methods that were developed to detect genes dysregulated in subsets of disease samples, such as COPA^[12]^, OS ^[13]^, ORT^[14]^ and MOST^[15]^. However, they are usually very sensitive to batch effects that, without corrections, may lead to false discoveries or to confound important subpopulation effects^[16]^. Prior application of normalization routines to the investigated samples are used to mitigate such technical biases, but improper normalization may still perturb the biological signal^[17]^. To tackle these difficult problems, Geman et al have firstly proposed to make use of the relative ordering information of gene expression within each sample, considering that the relative ordering of gene expression within each sample would be rather robust against batch effects and insensitive to data normalization^[18,19]^. The relative ordering of gene expression is overall stable in a particular type of normal human tissue but widely disturbed in diseased tissue. Zheng Guo et al has successfully developed several methods to detect patient specific differential expression information including mRNAs(RankComp)^[20]^, microRNAs (RankMiRNA)^[21]^, lncRNAs (LncRIndiv)^[22]^ and CpG sites^[23]^. These methods used pairs of molecules with a stable, relative order in a reference dataset to infer deregulated molecules in individual samples. Daniel Jost et al proposed PenDA^[24]^, which is based on the local ordering of gene expressions within individual cases and infers the deregulation status of genes in a sample of interest compared to a reference dataset. These efficient methodological tools allow to identify patient-specific molecular defects from the many precise molecular information and enable us to study cancer mechanisms in a novel way.

To date, none of methods above have been used in detecting differentially expressed proteins (DEPs) in individual patients. The relative ordering of protein expression in human normal and tumor tissues have not been investigated. There is a scarcity of systematic tool that could be used to evaluate the performance of individualized difference expression analysis algorithms in human cancer proteome. In this study, we reanalyzed the proteomic data of lung cancer, liver cancer, and gastric cancer, which included tumor and adjacent normal samples. These datasets shown that the within-sample relative expression orderings (REOs) of protein pairs in a particular type of normal tissue were indeed highly stable, which provided the basis for individual level DEPs analysis. Then, we evaluated five state-of-the-art tools (RankComp v1/v2, PENDA, Peng and Quantile) through classic computational (precision, Type one error control, parameter evaluation and robustness) and functional (pathway enrichment and survival analysis) criteria. A user-friendly tool kit, IDEP, were developed to enable implementation of personalized difference expression analysis algorithms.

## Material and methods

### Datasets

The following datasets were used in our experiments.

1. Lung-Xu dataset^[7]^: Protein abundance data of tumor (T) and tumor nearby tissues (TNT) in 103 patients with lung cancer
2. Lung-Gillette dataset^[9]^: Protein abundance data of T and TNT in 101 patients with lung cancer
3. Gastric-Ge dataset^[10]^: Protein abundance data of T and TNT in 84 patients with diffuse gastric cancer
4. Gastric-Ni dataset^[25]^: Protein abundance data of T and TNT in 54 patients with advanced gastric cancer
5. Liver-Gao dataset^[6]^: Protein abundance data of T and TNT in 159 patients with liver cancer

### Individualized differential expression analysis algorithms

The messages of the algorithm that we evaluated are listed below.

1. RankComp v1/v2 (https://github.com/pathint/reoa)
2. PENDA (https://github.com/bcm-uga/penda)
3. Peng method (https://github.com/xmuyulab/IDEPA/blob/master/deps_lib/methods_lib.py)
4. Quantile (https://github.com/xmuyulab/IDEPA/blob/master/deps_lib/methods_lib.py)
5. T-test (https://github.com/scipy/scipy)
6. Wilcoxon (https://github.com/scipy/scipy)

### Selection of reference group in individualized differential expression analysis algorithm

The determination of the reference group in the individualized differential expression analysis algorithm directly affects the performance of the algorithm. The reference group of RankComp v1 and RankComp v2 is to select the protein with stable expression order relative to the target protein in the normal cohort and tumor cohort. When RankComp v1 and RankComp v2 determine stable pairs, the former sets the threshold based on experience, and the latter sets the threshold based on the binomial distribution test. Penda selects proteins with stable expression order relative to the target protein and abundance close to the target protein from the normal cohort as the reference group. In the Peng method, the proteins in the normal cohort that have a stable abundance order relative to the target protein are first determined, and only the first three proteins with the smallest coefficient of variation are taken as the target protein reference group. However, Quantile uses the quantile method to determine the composition of the reference group. After determining the reference group, RankComp v1/v2 and PENDA determine the p value through statistical testing, and iterate repeatedly until the DE protein results are stable. The Quantile and Peng method determine the DEPs in the sample by setting the threshold.

### Overview of evaluation criteria

As shown in Figure 1, the evaluation of individualized difference expression analysis algorithm for protein abundance data is mainly divided into the following steps: preprocessing of protein abundance data; selection of protein reference group; individualized difference expression analysis; set evaluation indicators. The protein abundance data format was a two-dimensional matrix, the rows represented the names of the proteins, and the columns corresponded to the sample information. When setting evaluation indicators, the indicators are divided into two categories. The first category is the classic standard, including precision, Type one error control, impact of sample size, robustness and similarity. The second category is functional standards, including pathway enrichment analysis and survival analysis.

**Figure 1:**
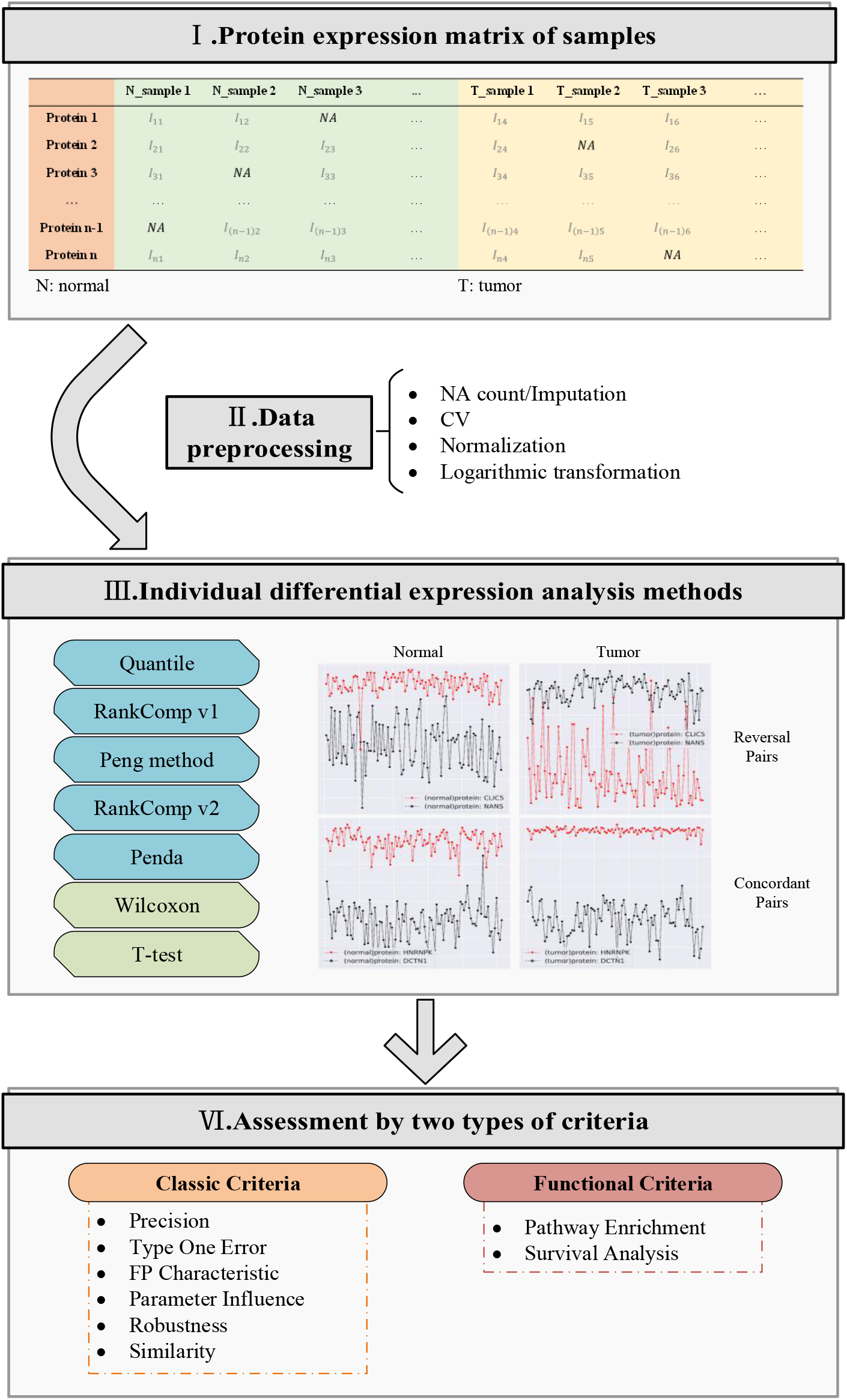
IDPE workflow.

### Identification of significantly stable REOs of protein pairs in normal tissues

We used REOA package (https://github.com/pathint/reoa) to identify significantly stable REOs of protein pairs in a type of normal tissue samples accumulated from different laboratories. For each protein pair (*P*_*i*_, *P*_*j*_), let n and k denote the total number of normal samples and the number of samples that have a certain REO pattern (*E*_*i*_>*E*_*j*_ or equally *E*_*j*_<*E*_*i*_), respectively, where *E*_*i*_ and *E*_*j*_ represent the expression abundance levels of *M*_*i*_ and *M*_*j*_, respectively. Then the probability of observing this REO pattern frequency (k/n) by chance is calculated by a binomial distribution test as following:

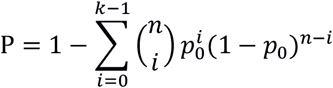

where *p*_0_ is 0.5, the probability of observing a certain REO pattern in a normal sample by chance. The P-values were adjusted using the Benjamini and Hochberg approach to control the FDR.

### Precision evaluation in real paired cancer-normal sample data

Five tumor proteome datasets (lung-Xu, lung-Gillette, gastric-Ge, gastric-Ni and liver-Gao) containing paired information were used to evaluate the precision of the difference expression analysis algorithms(RankComp v1/v2, PENDA, Peng method, Quantile, T-test and Wilcoxon signed-rank test). First, to meet the requirements of the T-test method for the normal distribution of data, the protein data were normalized and logarithmic., which did not affect the Rank-based algorithms. Second, 20 kinds of protein imputation methods in NAguideR^[26]^ were evaluated. Four criteria were used in the evaluation process, including Normalized root mean square error (NRMSE), NRMSE based sum of ranks (SOR), Average correlation coefficient between the original and imputed values (ACC/ACC OI), Procrustes statistical shape analysis (PSS). According to the evaluation results, BPCA is selected as the imputation method for protein abundance data. The maximum proportion of missing values is set to 30%, and data from different cohorts are filled separately. The difference information at the individual level is expected to be retained as much as possible, so there are no filter conditions for the coefficient of variation. Users can set the threshold of the coefficient of variation according to their requirements.

A part of the tumor data with paired information is selected as the gold standard data set to evaluate the algorithm precision, and the normal samples in the gold standard data were not used as the reference group. For example, in the lung-Xu data set with 103 pairs of paired samples, 51 pairs of samples (the normal sample group and the tumor sample group are represented as N_g_ and T_g_) were selected for the precision evaluation. The normal lung tissue sample group that did not contain N_g_ was used as the reference group to infer the protein differential expression status of the tumor sample group T_g_. We evaluated the precision of the differentially expressed (DE) protein identified for each cancer sample by evaluating whether the deregulation directions (upregulation or downregulation) of the DE protein predicted by the algorithm could be confirmed by the truly measured expression differences between the cancer sample and its adjacent normal tissue sample.

### Type one error control

Since there was no real tumor DE protein between samples of normal tissues, null data is constructed from normal tissue samples to evaluate the ability to control Type one error of algorithm. After the same preprocessing of precision evaluation, the protein abundance data of all normal tissue samples in the five data are extracted. To eliminate the differences introduced by different samples, the same null data contains normal sample information only from one dataset. And we generate null data with sample size of 6, 10, and 14, and the number of corresponding data sets are 40, 22, and 15 respectively. The individualized difference expression analysis method in IDPE was used to analyze the null data set to obtain the individualized difference expression results. P-value was set to 0.05, and the false discovery rate was calculated for each sample in the null data. We evaluated the type one error control by recording the fraction of tested proteins that were assigned a nominal P value of less than 0.05。

To compare the inherent deviations of different algorithms, according to the difference expression analysis results on the null data set, for each protein on each null data, we analyzed the characteristics of the False-positive proteins, including the average, variance, and coefficient of variation. Then we calculated the mean value of the protein characteristics on each null data and compare each of the three protein characteristics between DE and non-DE proteins ^[27]^. For each protein in each data set, we calculated the statistical value of the signal-to-noise ratio:

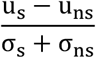

Where u_s_(u_ns_) and σ_s_(σ_ns_) respectively represent the mean and standard deviation of the characteristics on the DE protein (non-DE protein).

### The effect of the sample size of normal samples on the performance

In individualized difference expression analysis, normal tissue samples provide important reference for difference analysis of tumor tissue samples. Generally, the stability of the reference group increases as the number of normal tissue samples increases^[21]^.

In order to compare the effect of sample number in normal tissues on the algorithm performance, a subset of different sample numbers (20, 30, 40, 50, 60, 70, 80) is randomly selected from the normal tissue samples of lung-Xu data. We set the gold standard data for the subset according to the same method in precision evaluation, with a ratio of 0.5, and use the individualized difference expression algorithm in IDPE for analysis. According to the same method in precision evaluation, the gold standard data was constructed from the subset data, and the ratio was set to 0.5. The individual difference expression algorithm in IDPE was used for analysis. After obtaining the individualized difference analysis results, the precision and the number of difference proteins on the samples of the same subset are averaged. The same analysis was performed on the data sets gastric-Ge and liver-Gao. The number of generated samples by gastric-Ge is the same as the lung-Xu data set, and the number of generated samples in the liver-Gao dataset is 20, 40, 60, 80, 100, 120, 140.

### Robustness in individualized difference expression protein analysis

The robustness evaluation of the individualized difference expression algorithm focuses on the robustness of the method when it was applied to the same tumor tissue sample but the normal tissue sample was different. When there were real difference proteins in tumor tissue samples, no matter what normal tissue sample was used as the reference group, the real difference proteins should be able to be detected. Therefore, the difference expression results obtained from different reference groups for the same tumor tissue sample should be highly consistent. When the result consistency was not high, it means that the algorithm is obviously affected by the reference group, and the robustness of algorithm is low.

We sort the DE proteins according to their significance and select the area under the consistency curve^[27]^ of the top K proteins as the measuring standard of consistency. This area was also called the consistency area. More precisely, for each data set and corresponding individualized difference expression method, we sort the proteins in the sample by statistical significance (P-value or Q-value). For the individualized difference expression results obtained by two different reference sets, let k = 1, 2, ..., K, and we obtain the top k protein list of sample i in the two expression results. Then we calculated the number of shared proteins c_i_ for these two lists. The mean value c of c_i_ in all samples was calculated, and draw a consistency curve based on c and k. The area under the curve (consistency area) was used as the measuring standard of consistency. In order to obtain more interpretable values, we divide the consistency area by the largest possible value 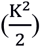. Therefore, when the normalized consistency area is 1, it means that the two individualized analysis results were the same. When the value was 0, it means that the two results are completely different. Compared with the simple intersection using the top K proteins, the consistency score contains the actual ranking of the top K proteins.

We used the normal tissue samples in the data sets lung-Xu and lung-Gillette as reference group 1 and reference group 2. The difference expression analysis algorithm was used to obtain the individualized difference analysis results of tumor samples in lung-Xu under these two reference groups. Based on the two difference expression results, the consistency score was calculated to compare the individual-level robustness of different algorithms on the lung data set. Similarly, the data sets gastric-Ni and gastric-Ge are also used to compare the individual-level robustness of different algorithms on the gastric data set.

To compare the group-level robustness of individualized difference expression analysis methods under different data sets in the same organization, seven difference expression analysis methods used to analyze lung-Xu and lung-Gillette (gastric-Ni and gastric-Ge). Then the individualized difference protein results were obtained. Through the binomial distribution test, the difference protein result at the group level can be obtained from the difference protein result at the individual level. The number of DE proteins in lung-Xu (gastric-Ni) was represented as N_1_, the number of DE proteins in lung-Gillette (gastric-Ge) was N_2_ and the number of proteins in the intersection of the DE proteins between lung-Xu (gastric-Ni) and lung-Gillette (gastric-Ge) is N_3_. The common rate can be expressed as:

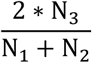

By comparing the common rate of different methods on different data sets, the group-level robustness of the individualized difference expression analysis method is obtained.

### Functional pathway enrichment

In order to further analyze the functions of proteins in tumor tissues, we obtained the DE proteins at the population level based on the results of individualized difference expression analysis and obtained significant pathways through pathway enrichment analysis. Finally, compare the significant paths obtained by RankComp v1/v2, Penda, Quantile, and Peng in different datasets.

Based on the binomial test, we obtained the results of group-level differential expression from the results of individualized differential expression analysis. Calculate the ratio p_de_ of deregulated proteins in the results of individualized differential expression in all samples relative to all proteins. Then judge whether the protein was differentially expressed at the group level according to the binomial test.

### Discovery of prognostic proteins

DE proteins may be related to patient prognosis, and individualized difference expression analysis tools provide individual-level difference expression information, which provides a new way for discovering treatment goals and personalized diagnosis. To compare the ability of different difference expression analysis tools to find prognostic proteins, the lung-Xu dataset and gastric-Ni dataset were analyzed based on the individualized difference expression analysis tool in IDPE (RankComp v1/v2, PENDA, Quantile, Peng method), and the results of individualized difference expression were obtained. In order to reflect the heterogeneity of tumors, for each method in each dataset, we retain proteins that are deregulated in 10%-90% of the samples. We grouped patients based on whether the protein was deregulated, and determined the prognostic protein by assessing whether there was a significant difference in the survival time of patients with or without DE protein. Use the univariate Cox proportional hazard regression model with 5% FDR control, we obtained the prognostic proteins found in different data sets through different methods, and obtain their survival curves.

## Results

### Highly stable REOs of proteins in normal tissues

The basic assumption of the individualized difference expression analysis algorithm is that protein pairs have stable relative expression orderings (REOs) in different normal samples. To verify whether there are stable pairs in the protein abundance data, we used lung cancer and gastric cancer as case studies. Table 1 shown statistical values of stable pairs in these two cancer types. There were 7,833,040 stable pairs of normal tissue samples in the lung-Xu data set, and 7,243,961 pairs in the lung-Gillette data set. 3,796,858 pairs were overlapped between these two datasets, which was much higher than the randomly. The percentage of overlapping stable protein pairs accounts for 48.47% for lung-Xu pairs. The stable pairs in the two data sets were highly consistent, reaching 99.91%. In the gastric cancer, the gastric-Ge data set and the gastric-Ni data set shown the same conclusions as the lung cancer. The POG in stomach tissue was as high as 90.59%, which is much higher than that in lung tissue. Fig. 2A shows the stable pairs of ANKFY1 and RPS3 proteins on the lung-Xu and lung-Gillette datasets. Figure 2B shows the stable pair of CORO1A and NBAS proteins on the gastric-Ge and gastric-Ni data sets. The abundance of ANKFY1 protein in the two different lung data sets was higher than that of RPS3. The CORO1A and NBAS proteins had the same performance on different gastric data sets.

**Table 1.** Reproducibility analysis of stable REOs of protein pairs in normal samples

| Tissue type | Group 1<sup>1</sup> | Group 2<sup>2</sup> | Random<sup>3</sup> | Overlap<sup>4</sup> | POG(%)<sup>5</sup> | Consistency(%)<sup>6</sup> |
| --- | --- | --- | --- | --- | --- | --- |
| Lung | 7,833,040 | 7,243,961 | 1,971,218 | 3,796,858 | 48.47 | 99.91 |
| Gastric | 4,166,089 | 4,068,172 | 1,908,348 | 3,774,167 | 90.59 | 99.97 |

1. Group 1 include Lung-Xu and Gastric-Ge dataset
2. Group 2 include Lung-Gillette dataset and Gastric-Ni dataset
3. Random denotes the expected number of overlapping stable protein pairs between Group 1 and Group 2 by chance with $P=0.05$ (hypergeometric test).
4. Overlap denotes the number of overlapping stable protein pairs between Group 1 and Group 2.
5. POG denotes the percentage of overlapping stable protein pairs among the list of stable protein pairs from Group 1.
6. Consistency denotes the percentage of overlapping stable protein pairs that had the same REO patterns in the two groups.

**Figure 2:**
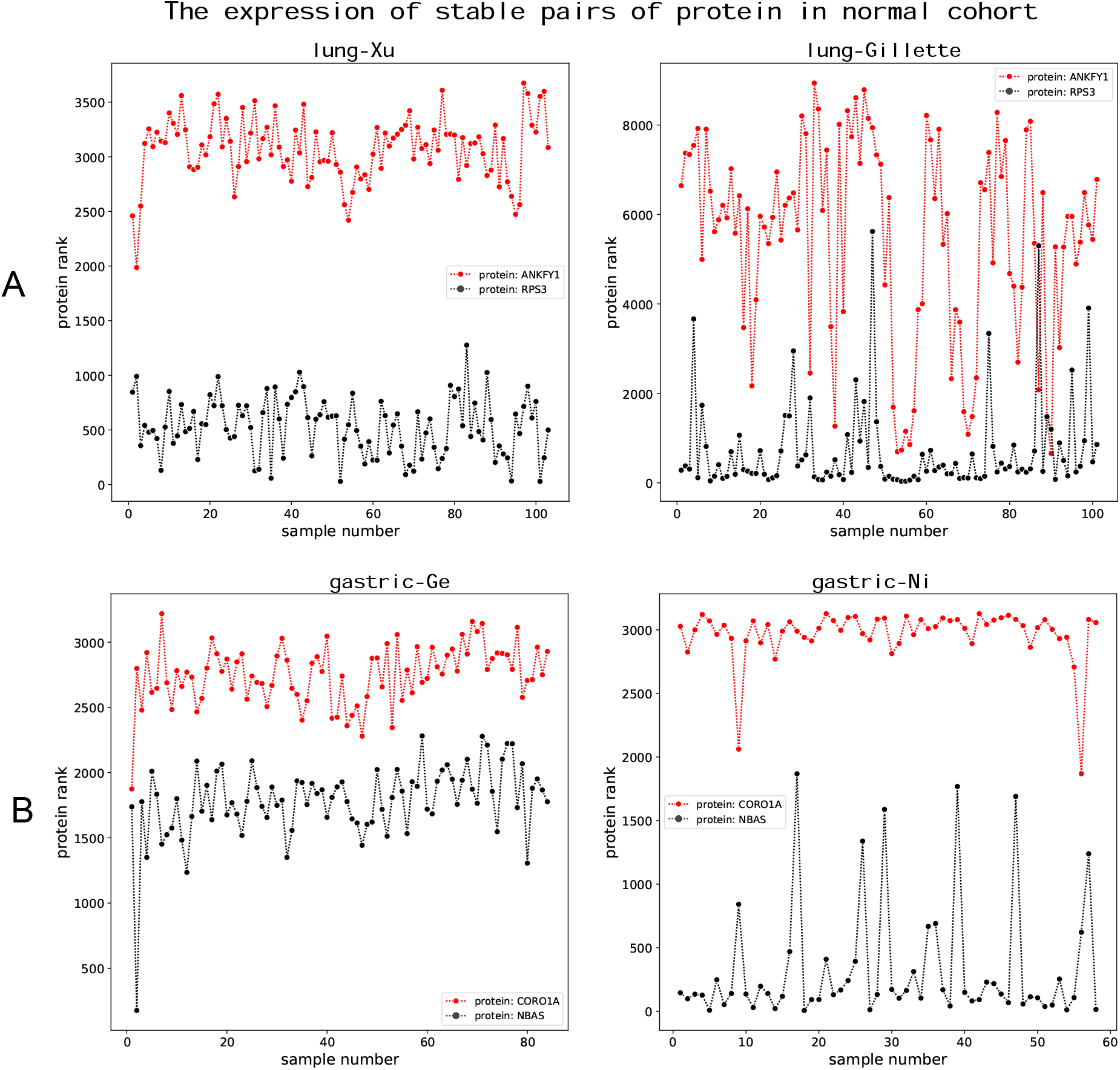
Stable protein pairs in different data sets. Each point in the figure represents the expression order of the protein in the sample. (A) The stable relationship between RPS3 and ANKFY1 on the data sets lung-Xu and lung-Gillette. (B) The stable relationship between CORO1A and NBAS on the data sets gastric-Ge and gastric-Ni.

These results above shown that, protein stable pairs had high consistency within the same cancer type. Protein stable pairs could be used as the reference group for individualized difference expression analysis methods to analyze the difference expression of proteins in tumor tissues at the individual level.

### Precision evaluation in real paired cancer-normal sample data

As shown in Fig. 3-1, the precisions of the individual-level difference expression analysis methods on the five independent datasets were much higher than that of the group-level methods, except for the Peng method. The Quantile method is from the perspective of outliers and the precision was the highest. The precision of PEND and RankComp v1 follow closely behind. The Peng method selects at most the top three proteins with the variation coefficient from small to large as the reference group. If the size of the reference group is too small, it will cause subtle changes in some proteins, which will seriously affect the difference protein results.

**Figure. 3–1:**
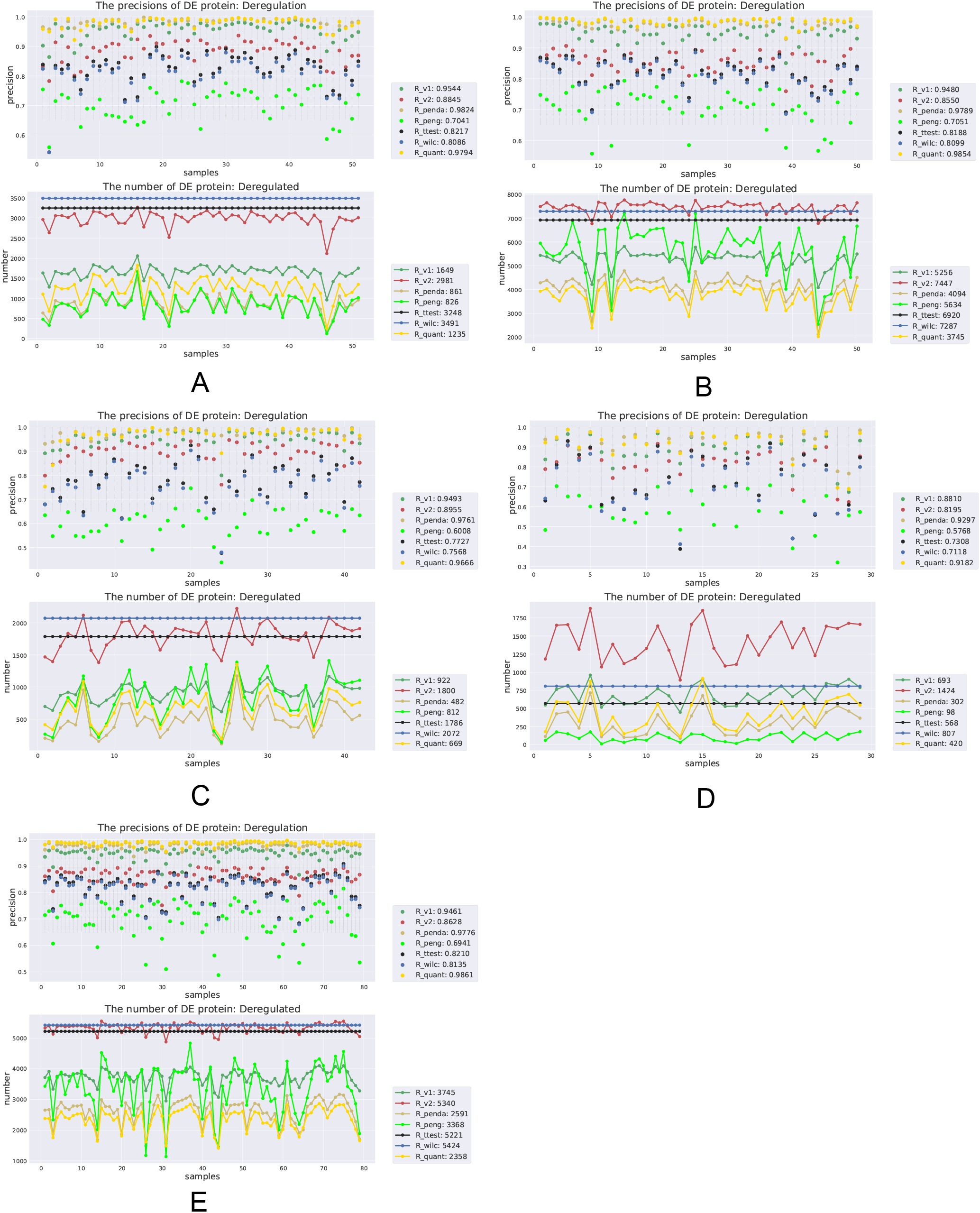
The precision and number of deregulated proteins on five datasets of seven differential expression analysis methods. (A) lung-Xu dataset. (B) gastric-Ge dataset (C) gastric-Ni dataset. (D) liver-Gao dataset. (E) lung-Gillette dataset.

From the point of view of the number of DE protein, the change trends of the seven methods were all consistent in the five independent datasets, indicating that the different methods had the consistent response direction to the changes in the number of the real difference data sample, but the sensitivities were different. In addition, the number of difference proteins found by RankComp v2 was the largest among the individual-level difference analysis methods. RankComp v2 used the binomial distribution test to select the reference group, and the FDR was 0.5. The simple cutoff point might result in high false discovery rates. Since RankComp v1/v2 has almost the same method in determining difference proteins, RankComp v2 performs lower precision than RankComp v1. Although both the T-test and Wilcoxon could find many difference proteins, the precision was not high. On different datasets, the difference protein data found by PENDA varies greatly. In the PENDA method, in addition to selecting the local stable pair as the reference group, the Quantile method was used as an alternative method, and method switching in PENDA was determined by data. Therefore, when analyzing different data sets, the difference protein data found varies greatly. The number of difference proteins by Peng method also varies greatly. The reason might be that the filter conditions in the Peng method were too strict, which causes many proteins to be excluded when determining the reference group, and will not enter the stage of difference protein analysis. In general, the Quantile and RankComp v1/v2 methods had relatively stable performance in terms of the precision and number of DEPs.

### Type one error control

Null data was used to evaluate the six methods on Type one error control, as shown in Fig. 3-2. The red line in the figure indicates that the theoretical FPR value of RankComp v1/v2, PENDA, T-test, and Wilcoxon is 0.05. Quantile considers outliers, so its theoretical FPR was 0. It could be seen from Fig. 3-2 A, B, and C that T-test and Wilcoxon were not sensitive to sample size in Type one error control. FPR of RankComp v1/v2 increased as the number of samples increases, while the FPR of PENDA and Quantile decreased as the number of samples increases. In terms of Type one error control, T-test and Wilcoxon perform best, indicating that the above two methods had small inherent deviations, while Penda and RankComp v2 had large inherent deviations.

**Figure. 3–2:**
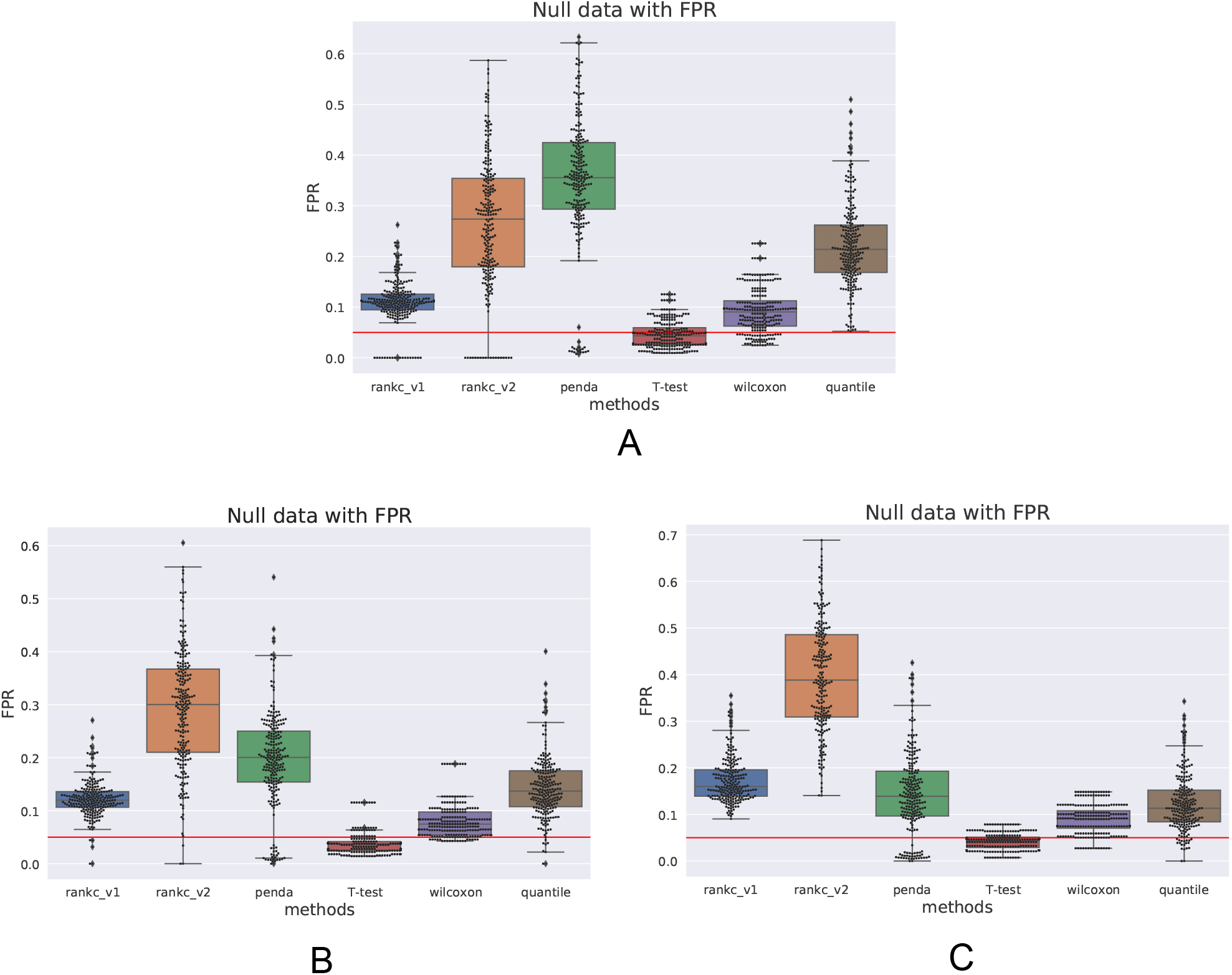
Type one error control of six differential expression analysis methods under null data of different sample sizes. (A) sample size = 6. (B) sample size = 10. (C) sample size = 14.

### The effect of the sample size of normal samples on the performance

When the normal samples were used as the reference group, we selected different normal sample subsets in the lung-Xu dataset, gastric-Ge dataset, and liver-Gao dataset as the reference groups. Different gradients were set according to the size of the data set. The influence of the number of normal samples on the performance of the difference expression analysis method was analyzed. As shown in Fig. 3-3, In the same differential expression analysis method, the precision of DE proteins remained stable in different numbers of normal samples, but the number of DE proteins changed differently.

**Figure. 3–3:**
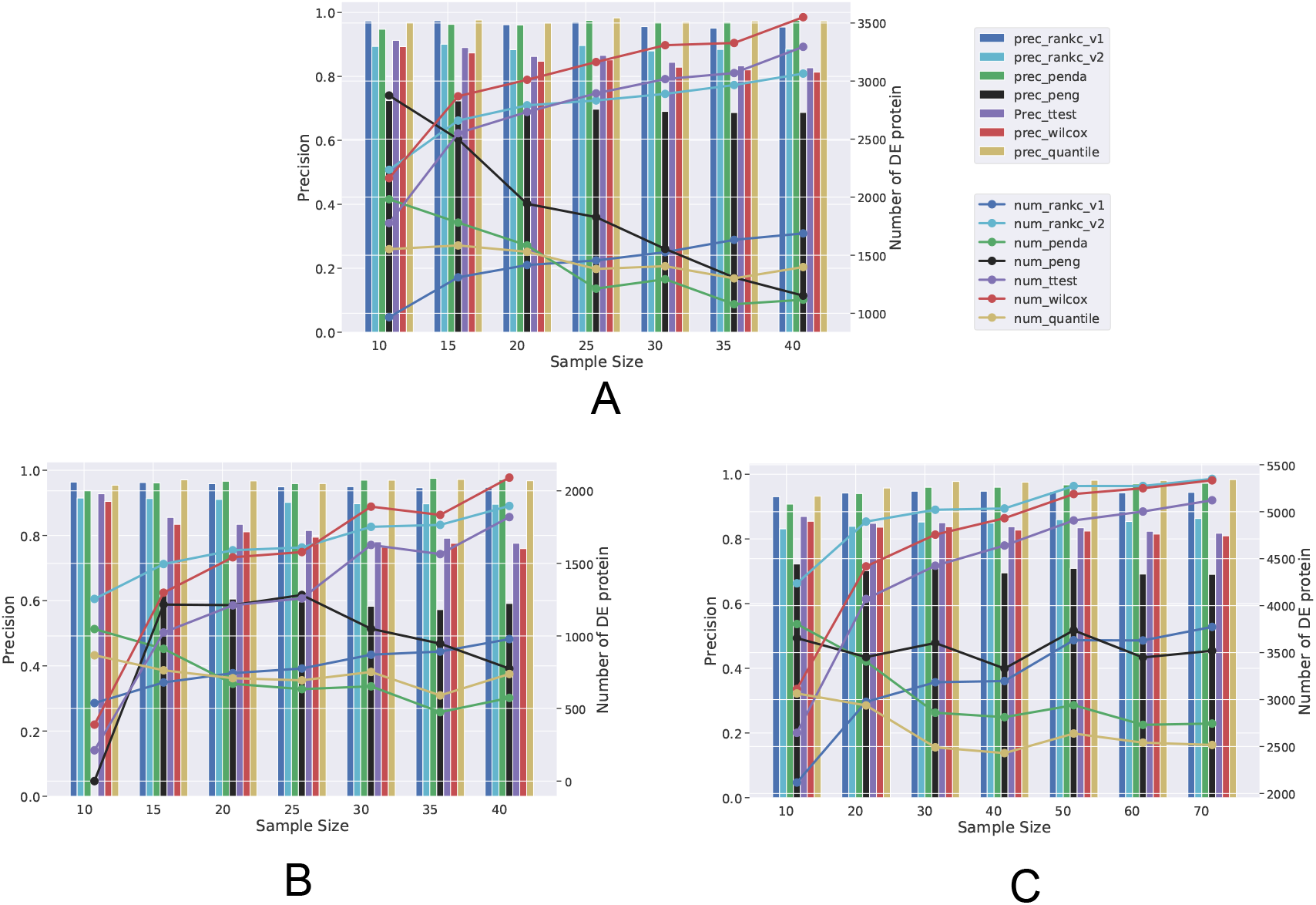
The effect of the sample size of normal samples in different datasets on the performance of seven differential expression analysis methods. (A) lung-Xu dataset. (B)gastric-Ge dataset. (C) liver-Gao dataset.

As the number of normal samples increases, the number of DE proteins found by RankComp v1/v2, T-test, and Wilcoxon also increased. T-test and Wilcoxon had a slight decrease in the precision of DE proteins, while the precision of the difference protein in RankComp v1/v2 remained stable. This may be due to the fact that T-test and Wilcoxon are group-level methods. The number of difference proteins in the PENDA and Quantile methods decreased with the increase of the number of normal samples, indicating that as the number of normal samples increases, the threshold in the method based on outliers became more stringent, resulting in a decrease in the number of difference proteins. For the Peng method, the changing trends of difference proteins were completely different in different data sets, indicating that the influence of the number of normal samples on the number of difference proteins was much smaller than the data set.

### Robustness in individualized difference expression protein analysis

We evaluated the robustness of these methods. As shown in Fig. 4 (A)-(D), the consistency scores of the difference expression analysis methods under normal samples in different data sets in the same tissue were compared at the individual level. DE proteins were subdivided into up-regulated proteins and down-regulated proteins. In lung tissues, difference expression analysis methods were inconsistent at the individual level of up-regulated proteins and down-regulated proteins. In the gastric organization, the performance was similar. Whether in up-regulated or down-regulated protein, RankComp v1/v2 was the most robust on the gastric data set. In terms of different protein types, the robustness of down-regulated proteins is lower than that of up-regulated proteins in lung cancer and gastric cancer.

**Figure 4:**
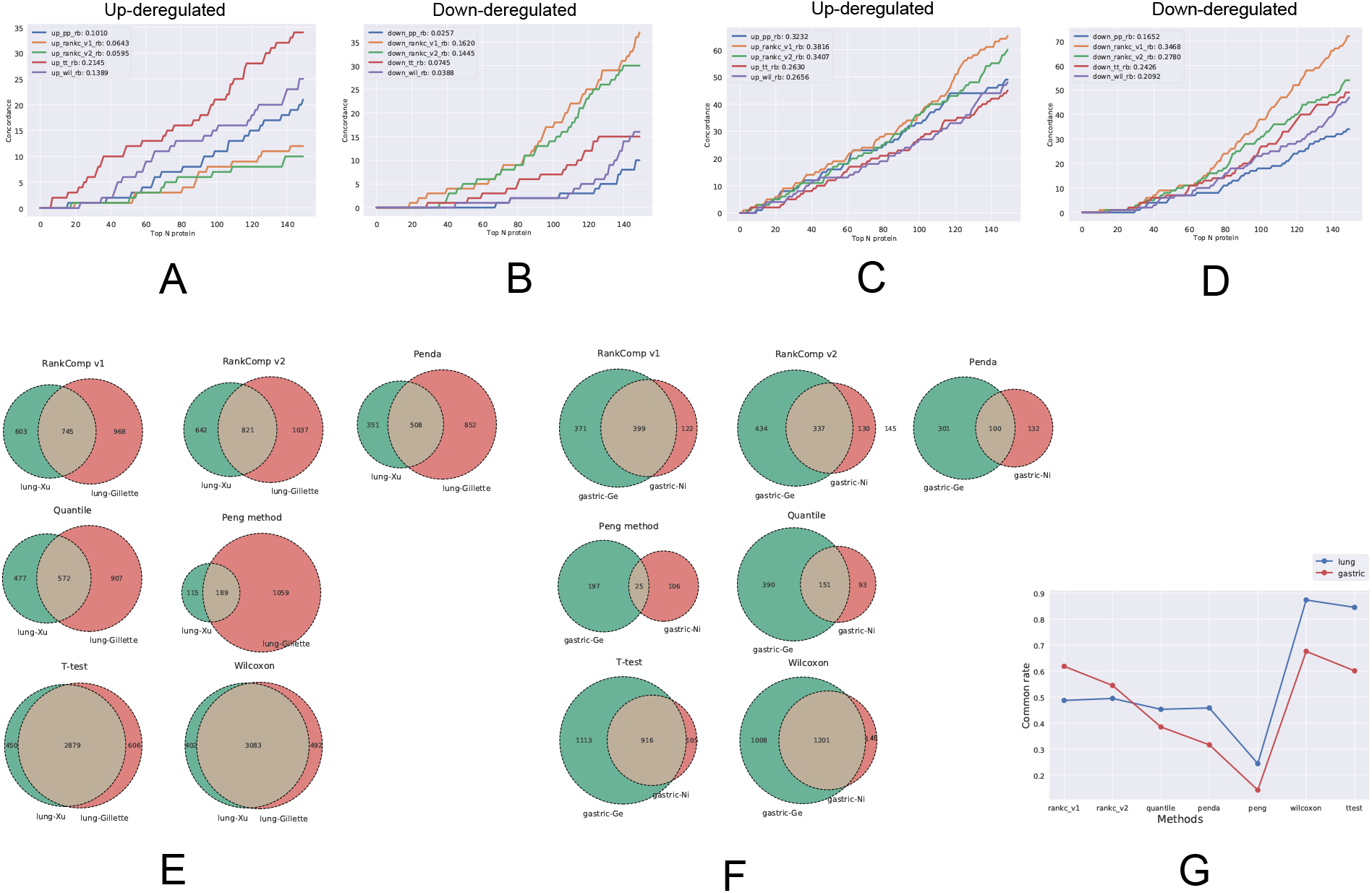
Robustness of differential expression analysis methods. (A-D) Evaluation of the individual-level robustness of five individualized differential expression analysis algorithms on the lung cancer dataset(A-B)and gastric cancer dataset(C-D). (E-F) Evaluation of the group-level robustness of seven individualized differential expression analysis algorithms on the lung cancer dataset(E) and gastric cancer dataset(F). (G) The common rates of seven individualized differential expression analysis algorithms on the lung cancer data and gastric cancer dataset.

As shown in Fig.4 (E)-(G), compared with individual-level methods, T-test and Wilcoxon could find more group-level difference proteins and had the highest common rates. For the individual level, RankComp v1 had the highest common rate, but the difference protein at the group level was less than RankComp v2. Among these methods, the Peng method relied largely on data sets and had the worst robustness.

### Functional pathway enrichment

In order to compare the differences in pathway functions determined by the individualized differential expression analysis method, we further analyzed the lung-Xu, lung-Gillette, gastric-Ge, and gastric-Ni data sets to obtain DE proteins at the population level, and used KEGG for enrichment analysis. We consider individual-level differential proteins that were differentially expressed on 10%-90% of the samples as group-level differential proteins. As shown in Figure 5, the performance of five methods in the data sets were inconsistent. In the lung data set, the significance pathways found by the individualized differential expression analysis method were highly consistent, while in the gastric data set, the pathways found by RankComp v1/v2, especially for metabolism pathways including Carbon metabolism, Propanoate metabolism and Pyruvate metabolism, were more significant and performed better. Due to the complexity and difference of biological data and functions, the results of pathway function enrichment analysis based on differential expression analysis were greatly affected by data sets. In general, compared to other methods, RankComp v1/v2 could find highly significant pathways in different tissues.

**Figure 5:**
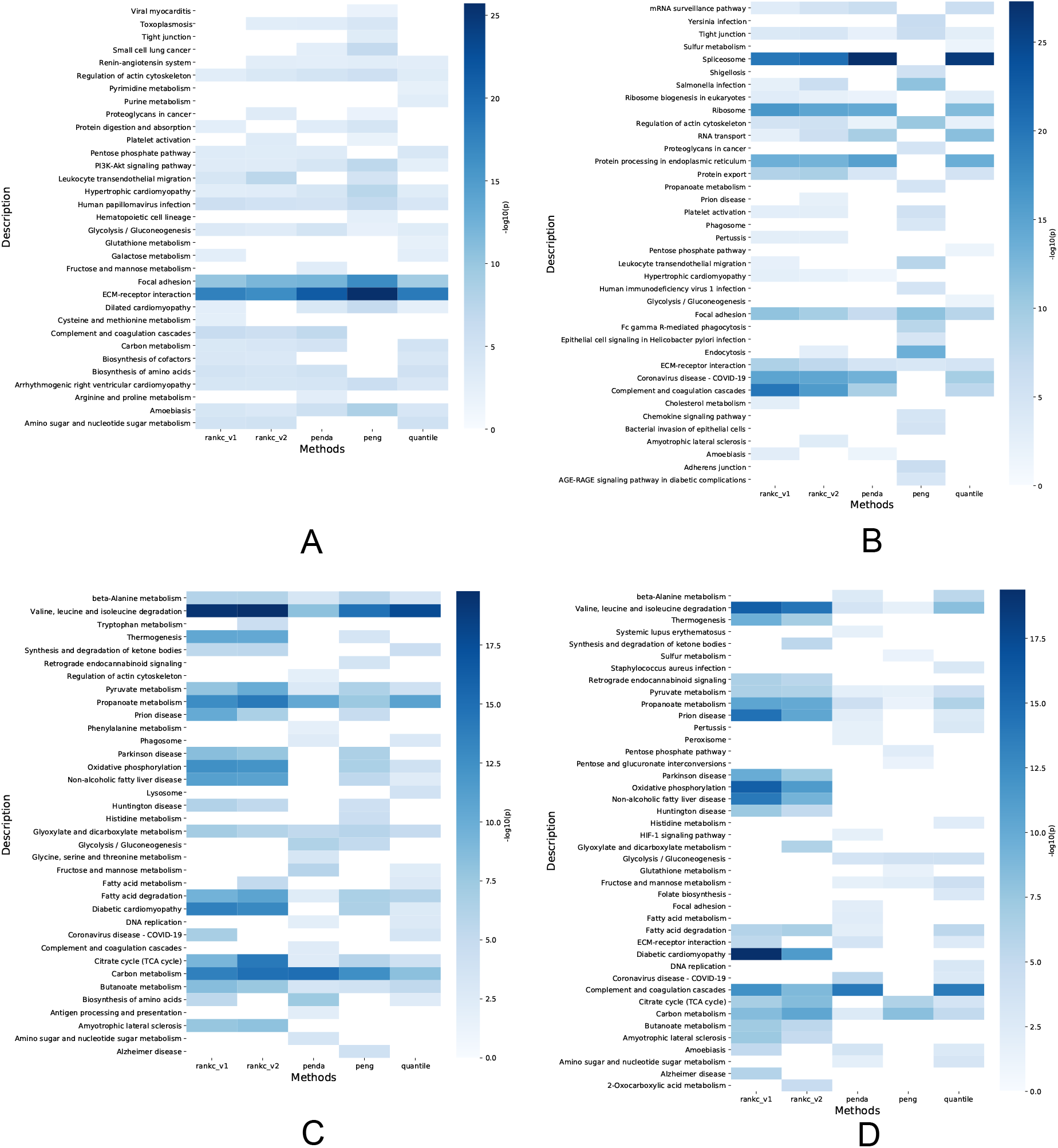
Functional pathway enrichment analysis of individualized differential expression analysis methods under different data sets. (A) lung-Xu dataset. (B) lung-Gillette dataset. (C) gastric-Ge dataset. (D) gastric-Ni dataset.

### Discovery of prognostic proteins

After obtaining the deregulated proteins, we further evaluated the prognostic values of DEPs from these individualized difference expression analysis methods using univariate cox proportional hazard regression model. We compared the survival analysis results including significance of cox model-based test and quantity of potential prognostic proteins. As shown in Fig. 6(A)(B), RankComp v1/v2 can find more potential prognostic proteins in both data sets and have a higher significance, and the number of prognostic proteins found by the other three methods is relatively close. Compared with the RankComp v2 method, RankComp v1 could find a higher proportion of potentially poor prognostic proteins. In addition, the data set also had a great influence on the number of prognostic proteins found by the individualized differential expression analysis method. The prognostic proteins found in RankComp v1, Quantile, Penda, and Peng in the gastric-Ni dataset were far less than those in the lung-Xu dataset. Here, we selected the EPS15 and HLA-DRA proteins in the lung-Xu data set and analyzed their survival curves. As shown in Fig. 6(C)(D), although different individualized difference expression methods have different ways of grouping samples, the significance of survival analysis results obtained were similar.

**Figure 6:**
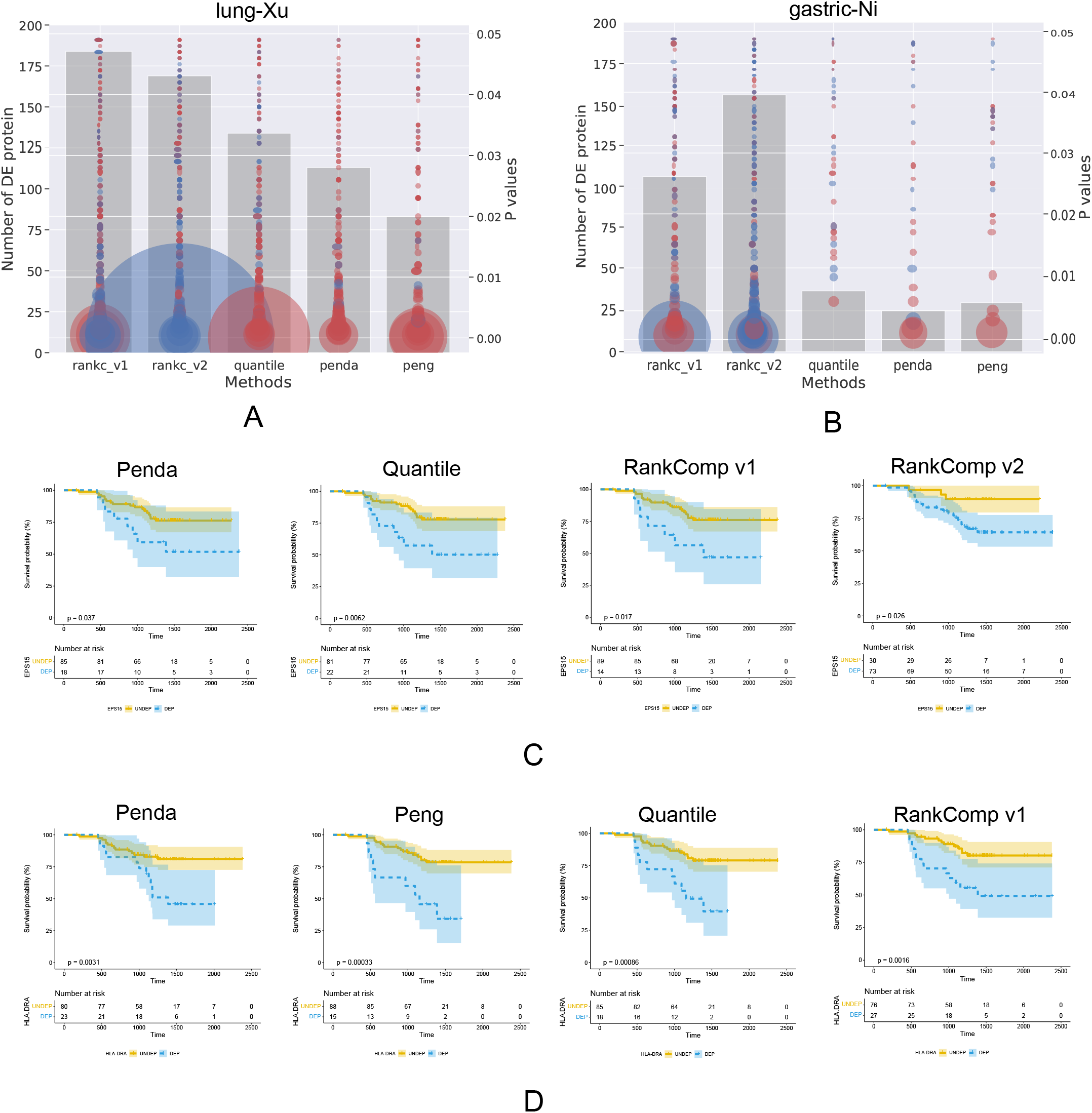
Prognostic proteins are found in five different individualized differential expression analysis methods in the lung-Xu dataset and the gastric-Ni dataset. A circle represents a protein, red represents a protein with HR>1, and blue represents a protein with HR<1. The area of the red circle corresponds to the absolute value of HR, and the area of the green circle corresponds to the reciprocal of HR. The height of the gray histogram corresponds to the number of prognostic proteins. (A) lung-Xu dataset. (B) gastric-Ni dataset. (C) The survival curve of EPS15 in the Lung-Xu dataset. (D) The survival curve of HLA-DRA in the Lung-Xu dataset.

## Discussion

The hopes of precision medicine rely on our capacity to measure individual molecular information for personalized diagnosis and treatment^[28]^. The differential expression analysis focusing on intergroup comparison can capture only DEPs at the population level, which may mask the heterogeneity of differential expression in individuals. Thus, to provide patient specific information for personalized medicine, it is necessary to conduct differential expression analysis at the individual level. Thanks to the development of efficient methodological tools that allow the identifications of molecular deregulation patterns at the individual level, getting better insights into inter-individual heterogeneities was made possible by the analyses of large cohorts of patients. This is the first study which systematically performed evaluation of personalized difference expression analysis algorithms in human cancer proteome. We found that these individualized difference analysis tools could reach much higher efficiency than the group-based methods. Using difference expression proteins between the tumor and adjacent normal samples as golden standard, we found the most frequently used population-level methods have larger false positive rates. Pathway enrichment results were dataset and analysis method dependent. RankComp v1/v2 could find highly significant metabolism pathways in different tissues.

The current IDEP has however several limitations. First, we used the difference expression proteins between the tumor and adjacent normal samples as golden standard. Such matched samples are usually rare which limit the evaluation in other cancer types. More general golden standard datasets and design are needed. Second, the datasets used here were mainly measured by the lable-free quantification methods. The within-sample REOs may be subject to a certain degree of uncertainty in samples measured by different platforms due to the differences in peptide quantification design principles. Thus, it is necessary to further evaluate the cross-platform properties of within sample REOs in order to extend the application scope of the individual-level differential expression analysis in human cancer proteome. Third, the current individual-level methods are not suitable for protein with low expression levels in all samples. More efficient filtering methods are needed to develop to improve the precision and robustness when applied in the proteome datasets.

To conclude, personalized differential analysis could provide a better understanding of disease-specific biological mechanisms and will promote the development of personalized therapeutic strategies in human cancer proteome.

We provide user guidelines so that IDEP could be run by users with limited computational experience. To ensure reproducibility of analysis, the penda vignette provides a summary of used parameters ready to be included in the method section of publications using IDEP.

## Notes

### Competing Interest Statement

The authors have declared no competing interest.

## Reference

[1] Rodriguez H, Zenklusen J C, Staudt L M, et al. The next horizon in precision oncology: Proteogenomics to inform cancer diagnosis and treatment[J]. Cell, 2021, 184(7): 1661–1670.

[2] Zhang B, Wang J, Wang X, et al. Proteogenomic characterization of human colon and rectal cancer[J]. Nature, 2014, 513(7518): 382–+.

[3] Zhang H, Liu T, Zhang Z, et al. Integrated Proteogenomic Characterization of Human High-Grade Serous Ovarian Cancer[J]. Cell, 2016, 166(3): 755–765.

[4] Mertins P, Mani D R, Ruggles K V, et al. Proteogenomics connects somatic mutations to signalling in breast cancer[J]. Nature, 2016, 534(7605): 55–+.

[5] Jiang Y, Sun A, Zhao Y, et al. Proteomics identifies new therapeutic targets of early-stage hepatocellular carcinoma[J]. Nature, 2019, 567(7747): 257–+.

[6] Gao Q, Zhu H, Dong L, et al. Integrated Proteogenomic Characterization of HBV-Related Hepatocellular Carcinoma[J]. Cell, 2019, 179(2): 561–+.

[7] Xu J-Y, Zhang C, Wang X, et al. Integrative Proteomic Characterization of Human Lung Adenocarcinoma[J]. Cell, 2020, 182(1): 245–+.

[8] Chen Y-J, Roumeliotis T I, Chang Y-H, et al. Proteogenomics of Non-smoking Lung Cancer in East Asia Delineates Molecular Signatures of Pathogenesis and Progression[J]. Cell, 2020, 182(1): 226–+.

[9] Gillette M A, Satpathy S, Cao S, et al. Proteogenomic Characterization Reveals Therapeutic Vulnerabilities in Lung Adenocarcinoma[J]. Cell, 2020, 182(1): 200–+.

[10] Ge S, Xia X, Ding C, et al. A proteomic landscape of diffuse-type gastric cancer[J]. Nature Communications, 2018, 9.

[11] Tusher V G, Tibshirani R, Chu G. Significance analysis of microarrays applied to the ionizing radiation response[J]. Proceedings of the National Academy of Sciences of the United States of America, 2001, 98(9): 5116–5121.

[12] Tomlins S A, Rhodes D R, Perner S, et al. Recurrent fusion of TMPRSS2 and ETS transcription factor genes in prostate cancer[J]. Science, 2005, 310(5748): 644–648.

[13] Tibshirani R, Hastie T. Outlier sums for differential gene expression analysis[J]. Biostatistics, 2007, 8(1): 2–8.

[14] Wu B. Cancer outlier differential gene expression detection[J]. Biostatistics, 2007, 8(3): 566–575.

[15] Lian H. MOST: detecting cancer differential gene expression[J]. Biostatistics, 2008, 9(3): 411–418.

[16] Leek J T, Scharpf R B, Bravo H C, et al. Tackling the widespread and critical impact of batch effects in high-throughput data[J]. Nature Reviews Genetics, 2010, 11(10): 733–739.

[17] Wang D, Cheng L, Wang M, et al. Extensive increase of microarray signals in cancers calls for novel normalization assumptions[J]. Computational Biology and Chemistry, 2011, 35(3): 126–130.

[18] Geman D, D’avignon C, Naiman D Q, et al. Classifying gene expression profiles from pairwise mRNA comparisons[J]. Statistical applications in genetics and molecular biology, 2004, 3: Article19–Article19.

[19] Tan A C, Naiman D Q, Xu L, et al. Simple decision rules for classifying human cancers from gene expression profiles[J]. Bioinformatics, 2005, 21(20): 3896–3904.

[20] Wang H, Sun Q, Zhao W, et al. Individual-level analysis of differential expression of genes and pathways for personalized medicine[J]. Bioinformatics, 2015, 31(1): 62–68.

[21] Yan H, Cai H, Guan Q, et al. Individualized analysis of differentially expressed miRNAs with application to the identification of miRNAs deregulated commonly in lung cancer tissues[J]. Briefings in Bioinformatics, 2018, 19(5): 793–802.

[22] Peng F, Wang R, Zhang Y, et al. Differential expression analysis at the individual level reveals a lncRNA prognostic signature for lung adenocarcinoma[J]. Molecular Cancer, 2017, 16.

[23] Yan H, He J, Guan Q, et al. Identifying CpG sites with different differential methylation frequencies in colorectal cancer tissues based on individualized differential methylation analysis[J]. Oncotarget, 2017, 8(29): 47356–47364.

[24] Richard M, Decamps C, Chuffart F, et al. PenDA, a rank-based method for personalized differential analysis: Application to lung cancer[J]. Plos Computational Biology, 2020, 16(5).

[25] Ni X, Tan Z, Ding C, et al. A region-resolved mucosa proteome of the human stomach[J]. Nature Communications, 2019, 10.

[26] Wang S, Li W, Hu L, et al. NAguideR: performing and prioritizing missing value imputations for consistent bottom-up proteomic analyses[J]. Nucleic Acids Research, 2020, 48(14).

[27] Soneson C, Robinson M D. Bias, robustness and scalability in single-cell differential expression analysis[J]. Nature Methods, 2018, 15(4): 255–+.

[28] Vitali F, Li Q, Schissler A G, et al. Developing a “personalome’ for precision medicine: emerging methods that compute interpretable effect sizes from single-subject transcriptomes[J]. Briefings in Bioinformatics, 2019, 20(3): 789–805.

